# Secreted Protein Production is Improved by Controlling Endoplasmic Reticulum Stress Associated Protein Degradation

**DOI:** 10.1101/2025.08.05.666879

**Authors:** R. Chauncey Splichal, Christina Chan, S. Patrick Walton

## Abstract

Graphical Abstract

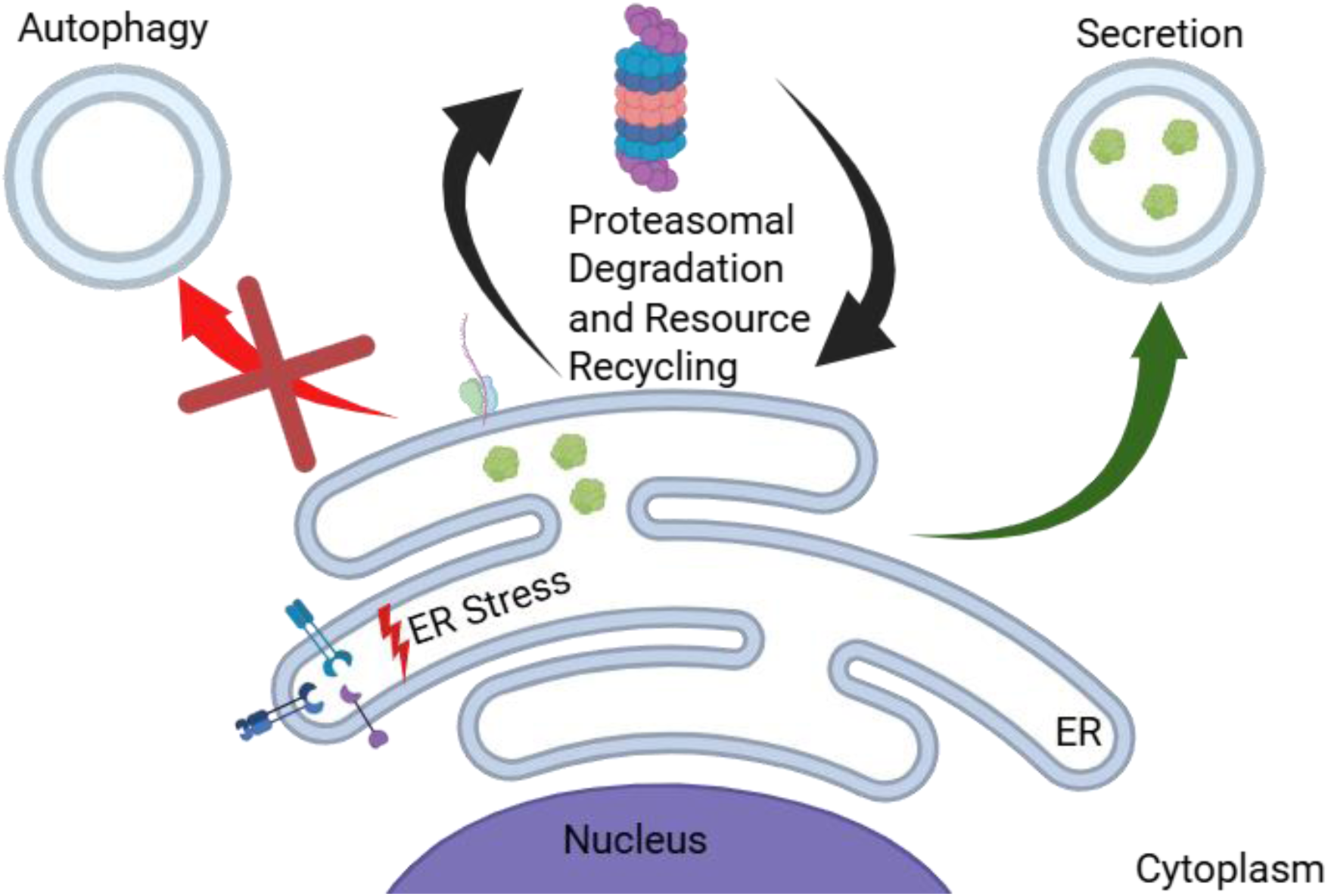

Therapeutic proteins are produced frequently by mammalian cells in large-scale bioreactors. As a result, producer cells are exposed to a chemically (nutrients, gas exchange, target protein overexpression) and physically (shear due to mixing) stressful environment, which can lead to loss of proteostasis and endoplasmic reticulum (ER) stress. In response, cells activate the unfolded protein response (UPR). The UPR includes activation of autophagy and proteasomes, both of which target unfolded/misfolded proteins for degradation. To investigate the impacts of autophagy and proteasome activity on secreted protein production in ER-stressed cells, we used HeLa and MDA-MB-231 cells transfected to express *Gaussia* luciferase (as a model for therapeutic protein production) and exposed to tunicamycin (TM) (to activate ER stress). As expected, TM exposure decreased protein production and secretion. Inhibiting autophagy improved secretion in stressed cells as expected. However, counterintuitively, increasing proteasomal degradation improved secretion while inhibiting proteasomal activity decreased secretion, that is proteasomal activity was directly correlated to secretion. Taken together, our results demonstrate that protein secretion can be improved through control of autophagy and proteasomal activity, providing insight into strategies for improving yield from protein production bioprocesses.

**Key Points:**

1. Tunicamycin induced ER stress reduced protein production.
2. Autophagy inhibition improved secretion in ER stressed cells.
3. Activation of proteasomal degradation improved secretion in ER stressed cells.

## Introduction

Recombinant proteins have a wide range of uses in medicine from diagnosis and prevention of diseases to treatments for cancer and infectious diseases. Industrial cell culture of mammalian cells requires the use of temperature-controlled bioreactors with mixing of highly viscous cultures. Agitation causes shear stress and nonetheless leaves heterogeneities in temperature, nutrients, and oxygen (Kaufmann et al. 1999; Keane et al. 2003; Handlogten et al. 2018; Coulet et al. 2022). In response to these kinds of environmental stressors, cells, especially those overexpressing a product protein, can experience a loss of proteostasis, causing the protein quality control machinery in the endoplasmic reticulum (ER) to become overwhelmed, a condition referred to as ER stress(Schröder and Kaufman 2005; Hetz et al. 2020; Splichal et al. 2024). While steps can be taken to mitigate ER stress in bioreactors, due to the requirements of industrial bioprocesses, cells are always experiencing ER stress(Godoy-Silva et al. 2009; Ha et al. 2019; Sinharoy et al. 2020).

Cells respond to ER stress via the unfolded protein response (UPR). The UPR has three pathways mediated by three distinct sensor proteins: Inositol-Requiring Enzyme 1α (IRE1α),(Schröder and Kaufman 2005) Protein Kinase R-like Endoplasmic Reticulum Kinase (PERK), and Activating Transcription Factor 6 (ATF6)(Hetz 2012; Splichal et al. 2024). IRE1α dimerizes and autophosphorylates in the presence of misfolded proteins, activating an RNase domain that cleaves X-box Binding Protein-1 (XBP1) mRNA into the XBP1-spliced (XBP1-s) mRNA, which encodes for the transcription factor XBP1-s(Hetz 2012). XBP1-s initiates signaling that leads to an increase in the size of the ER, upregulates chaperone protein synthesis, and activates ER-associated degradation (ERAD) of excess/misfolded proteins(Hwang and Qi 2018). PERK dimerizes in response to ER stress and activates Activating Transcription Factor 4 (ATF4) to increase chaperone production and C/EBP homologous protein (CHOP) pathways(Diehl and McQuiston 2017). PERK also phosphorylates eukaryotic translation initiating factor 2α (eIF2α), which results in an overall reduction of protein synthesis by reducing mRNA incorporation into ribosomes(Diehl and McQuiston 2017). ER stress causes dissociation of ATF6 from Binding Immunoglobulin Protein (BiP, GRP78, HSPA5). In turn, ATF6 translocates to the Golgi apparatus where it is cleaved by Site-1 protease (S1P) and Site-2 protease (S2P) to form the nuclear factor ATF6(Lei et al. 2024). Nuclear factor ATF6 amplifies the IRE1α and PERK responses to ER stress by upregulating XBP1 mRNA expression, chaperone expression, and CHOP expression. Overall, the UPR upregulates chaperone protein synthesis, decreases global protein production, increases protein degradation of misfolded or unfolded proteins, and, if proteostasis is not restored after these measures, the UPR induces apoptosis via the PERK-eIF2a-ATF4-CHOP pathway(Hetz 2012; Splichal et al. 2024).

The UPR also alters cellular secretion pathways. The endomembrane system connects the ER, Golgi, plasma membrane, and lysosomes through membrane bound vesicles. XBP1-s activation enlarges the size of the ER and ER-associated vesicles to reduce the concentration of misfolded proteins(Shaffer et al. 2004; Tigges and Fussenegger 2006). The UPR promotes protein degradation through ERAD and autophagy, and both degrade proteins that could otherwise be secreted(Meusser et al. 2005; Ichimiya et al. 2020). ERAD occurs through enhancement of proteasomal degradation in three steps: recognition, extraction, and degradation(Meusser et al. 2005). Misfolded proteins are recognized by chaperone proteins such as heat shock proteins ER DnaJ 3 (ERdj3), ERdj4, and ERdj5(Daverkausen-Fischer et al. 2022). Retrotranslocation proteins recognize and bind misfolded proteins in the ER lumen and undergo a conformational change to move the misfolded protein to the cytoplasm, where they are ubiquitinated by UPR regulated E3 enzymes including Suppressor Enhancer Lin-12-Like (SEL1L), Ubiquitin Protein Ligase E3 Component N-Recognin 4 (UBR4), and Ubiquitin Protein Ligase E3 Component N-Recognin 5 (UBR5)(Tang et al. 2020; Lin et al. 2024). Ubiquitinated proteins are trafficked to the 26S proteasome and degraded(Meusser et al. 2005). Because ERAD only degrades soluble proteins(Meusser et al. 2005; Fujita et al. 2007), cells need additional pathways for degradation of insoluble proteins and other cellular components.

Autophagy or “self-eating” refers to three different processes: microautophagy, chaperone-mediated autophagy (CMA), or macroautophagy(Ichimiya et al. 2020). Microautophagy is the homeostatic lysosomal degradation of material and uses direct transport of cargo into lysosomes(Wang et al. 2023). In CMA, heat shock protein (HSP)A8 binds misfolded proteins for retrotranslocation to the cytoplasm and subsequent insertion of the misfolded protein into the lysosome for degradation(Dice 2007). Most commonly, as we do here, autophagy refers to macroautophagy. Macroautophagy breaks down insoluble material or large amounts of material to maintain cellular function; macroautophagy is active in degrading protein aggregates, recycling organelles, and generating substrates and intermediates in response to nutrient deprivation(Yu et al. 2018). In macroautophagy, Unc-51-like Kinase 1 (ULK1) initiates a membrane formation event that creates a phagophore to encapsulate material to be degraded(Petherick et al. 2015). This vesicle matures into a double membrane vesicle called an autophagosome. Autophagosomes merge with lysosomes to initiate degradation of their contents(Ichimiya et al. 2020). Thus, ERAD and autophagy, which are both activated by ER stress, result in altered trafficking of expressed proteins (i.e., to degradation rather than secretion).

Here, we modeled ER stress on protein production in cell culture, with the goal of understanding how ER stress alters protein trafficking and secretion. Our results demonstrated that the UPR in response to ER stress (specifically IRE1α activation) improved protein secretion. Importantly, we show that, in cells under ER stress, inhibiting autophagy and enhancing proteasomal degradation improved secretion of proteins, presenting a potential opportunity for improving therapeutic protein secretion in industrial bioprocesses.

## Methods and Materials

### Cell Culture

HeLa and MDA-MB-231 cells were purchased from ATCC. All cells were maintained in high glucose DMEM (Gibco 11965092) containing 10% FBS (R&D Systems S11550) and 50 mM HEPES (Fisher BP410-500) at 5% CO_2_ and 37C. Cells were passaged three times per week. MDA-MB-231 IRE1α knockout cells were generated using CRISPR-Cas9 knockout with plasmids gifted by Dr. Feng Zhang (Addgene plasmid # 48138 and # 62988) and Dr. Andrea Ventura (Addgene plasmid # 64073).

### Nucleic Acid Preparation

A plasmid for expressing secreted *Gaussia princeps* luciferase was generously provided by Dr. R. Alexander Wesselhoeft and Dr. Daniel G. Anderson. Plasmid was linearized using Xba1 restriction digest enzyme (NEB R0145S) per manufacturer’s instructions prior to in vitro transcription according to manufacturer’s instruction using T7 high yield *in vitro* transcription kit (Thermo K0441) and MEGAClear transcription clean-up kit (Thermo AM1908). Plasmid DNA coding for secreted *Gaussia* luciferase conjugated to cyan fluorescent protein was generously provided by Professor Andrew Dillin at UC Berkeley. Bacteria were grown in LB broth (Affymetrix 75852) containing 200 µg/mL ampicillin (Fisher BP1760-25). Plasmid DNA was isolated using a ThermoFisher Gene Jet Plasmid Miniprep (K0503).

### Transfections

All cells were plated in 0.5 mL of DMEM containing 80,000 cells/mL in a 24 well plate (Costar 3526). 24 hours later, culture DMEM was replaced with serum free DMEM 1 hour prior to transfection. Nucleic acids were mixed with LF2K (Invitrogen 52887) per manufacturer’s instructions. Final concentrations (once added to the cells) for RNA transfections were 160 ng/mL RNA and 200 ng/mL LF2K and for pDNA transfection final concentrations were 200 ng/mL and 1200 ng/mL LF2K. Transfection solutions were added to cells in serum free DMEM and allowed to incubate at 37C for 4 hours. After cells were washed with 1 mL culture DMEM, fresh culture DMEM containing stressors, inhibitors, or activators was added, and cells recovered for 24 hours before analysis. Concentrations of chemicals used were TM (T7765-5MG) – 5 µg/mL 3MA (SAE0107-10ML) – 5 mM Rolipram (Bio-Techne 0905)– 60 µM VR23 (AdipoGen 1624602-30-7) – 3 µM.

### Luciferase Measurement

Luciferase measurements were performed using the Thermofisher *Gaussia* Luciferase Kit (Thermo 16161). 20 µL of recovery media was added a well in a Corning 3904 black wall 96 well plate. The remaining media was aspirated, and cells were washed with 1 mL of DPBS. Cells were lysed in 150 µL of 2X cell lysis solution mixed with 150 µL of DPBS. Plates containing lysis solution were placed on an orbital shaker at 65 rpm for 1 hour. 50 µL of lysis solution was added to the 96-well plate. For each test, 0.5 µL of 100X luciferase reagent solution was mixed with 49.5 µL of DPBS. The plate was loaded into a BioTek Synergy H1 Microplate Reader and shaken for 25 seconds, then raw luminescence was collected for 3 seconds in each well.

### Microscopy

Cells were plated Millicell EZ Slides (PEZGS0816) at 90000 cells/mL in 500 µL of culture DMEM. Cells were treated with the same conditions described above. After treatment, cells were fixed by adding 500 µL of 3:1 Methanol (Sigma 34860-4L-R): Glacial Acetic acid (Sigma AX00073-75). After 5 minutes, the culture media/fixative mixture was removed and replaced with 500 µL of fresh 3:1 Methanol:Glacial Acetic Acid for 10 min. Cells were rinsed for 5 min in DI water twice. LC3B staining was for 8 hours at 1:500 LC3B-DyLight550 antibody (Novus NB100-2220R): DPBS. Fixed and stained cells were rinsed three times with fresh DPBS before 40 µL of Prolong Antifade Glass Mount (Invitrogen P36984) was added to each well surface and a glass coverslip was applied. The mount was allowed to cure overnight before imaging on the Leica Stellaris 5 Confocal Laser Scanning microscope at 100x. LC3B-DyLight was imaged at Ex:562nm/Em 572-700nm. CFP was imaged at Ex 488nm/Em 500-550nm.

### Statistics

Statistics were calculated first using two-way ANOVA to determine significance followed by Student’s t-test to calculate pair-wise p-values. Measurements were made using 3 plates of cells with 3 replicates using independent solutions for each plate.

## Results

### ER stress reduces secretion relative to intracellular accumulation

To investigate the impact of ER stress on the secretion of expressed proteins, we tested the ability of HeLa and MDA-MB-231 cells to synthesize and secrete *Gaussia princeps* luciferase (Gluc) from transfected mRNA in the presence of increasing concentrations of the N-glycosylation inhibitor tunicamycin (TM) (Figure 1). Inhibition of N-glycosylation by TM reduces conformational stability and increases aggregation, leading to accumulation of misfolded proteins and ER stress(Jayaprakash and Surolia 2017). Increasing concentrations of TM reduced cell numbers for both HeLa and MDA-MB-231 cells (Figure 1A), likely due to both reduced growth rate and cytotoxicity. Increasing TM concentration also reduced the amount of Gluc secreted into the culture media (Figure 1B). Plotting cell number vs. Gluc secreted showed a significant correlation (Figure 1C). In contrast, increasing levels of TM resulted in no change (HeLa) but increased (MDA-MB-231) Gluc in cell lysates (Figure 1D). Lysate Gluc signal and cell number were not correlated (Figure 1E). To capture both these results simultaneously, we established a metric for the efficiency by which cells secrete expressed protein, the ***secretion ratio***, which is the quantity of Gluc in the media divided by the quantity in the cell lysate.

**Figure 1.**
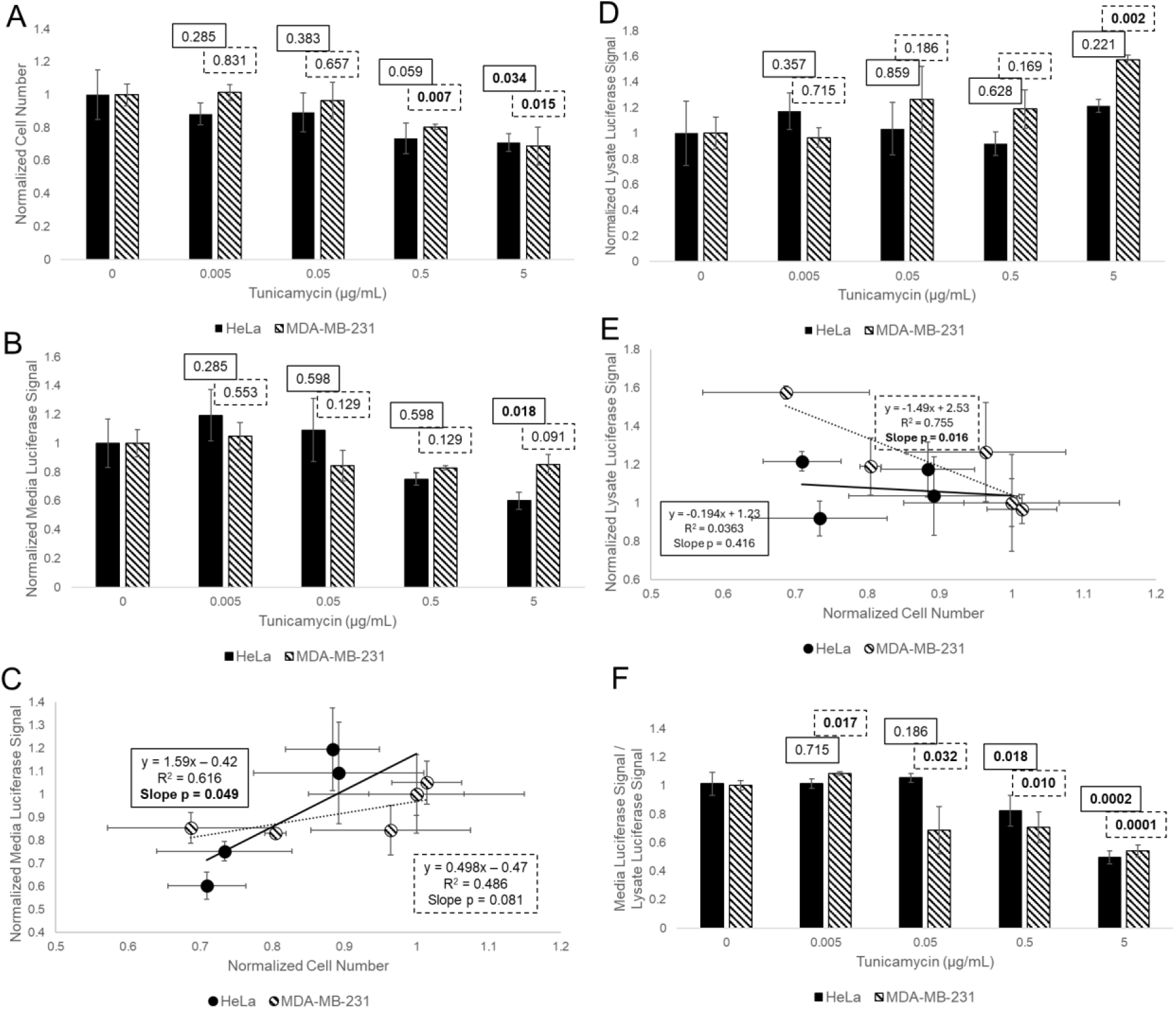
The impact of tunicamycin treatment on protein secretion. A) Normalized cell number vs. TM concentration for HeLa (solid) and MDA-MB-231 (striped) cells. B) Normalized 24-hour Gluc accumulation in the media vs TM concentration. C) Normalized cell number vs. normalized 24-hour media Gluc accumulation. D) Normalized Gluc in the cell lysate 24-hours post mRNA transfection vs. TM concentration. E) Normalized cell number vs. normalized Gluc in cell lysate. F) Normalized secretion ratio (24-hour media accumulation of Gluc divided by normalized Gluc in cell lysate) vs. TM concentration. *Error bars represent standard deviations. Values indicate Student’s t-test pair-wise p-values compared to 0 µg/mL TM, N = 3. Bold indicates p < 0.05*.

Secretion ratio decreased with increasing TM for both cell lines (Figure 1F). The changes in secretion ratio were associated with activation of autophagy (see supplemental figures 1A-F for representative micrographs of Gluc retention and the autophagosome marker Microtubule-associated proteins 1A/1B light chain 3B (LC3B)). Taken together these results suggest that ER stress i) reduced overall Gluc production in both cell lines by limiting cell growth, and ii) increased the fraction of the expressed, functional Gluc that was sequestered within the cells rather than secreted.

### IRE1α mitigates the impact of ER stress on secretion

IRE1α splicing of XBP1 mRNA and subsequent transcriptional changes are known to be important cellular responses to maintain secretory capacity(Ku et al. 2010). To investigate the role of IRE1α activity on secretion under ER stress, we used MDA-MB-231 wild type (WT) and IRE1α^-/-^ (KO) cells. TM treatment decreased the cell number and media accumulation of Gluc in both the WT and KO cells (Figures 2A&B). The amount of Gluc that accumulated in WT TM-treated cells was similar to the amount in WT cells (Figure 2C, left two columns). In contrast, the amount of Gluc that accumulated in the TM-treated KO cells was greater than the amount that accumulated in untreated KO cells (Figure 2C, right two columns). Thus, knocking out IRE1α further reduced the secretion ratio in response to ER stress induced by TM (Figure 2D, see supplemental figure 2 for representative micrographs depicting increased accumulation and autophagy maker LC3B in KO cells). Hence, IRE1α function is important for supporting secretion in cells experiencing TM-mediated ER stress.

**Figure 2.**
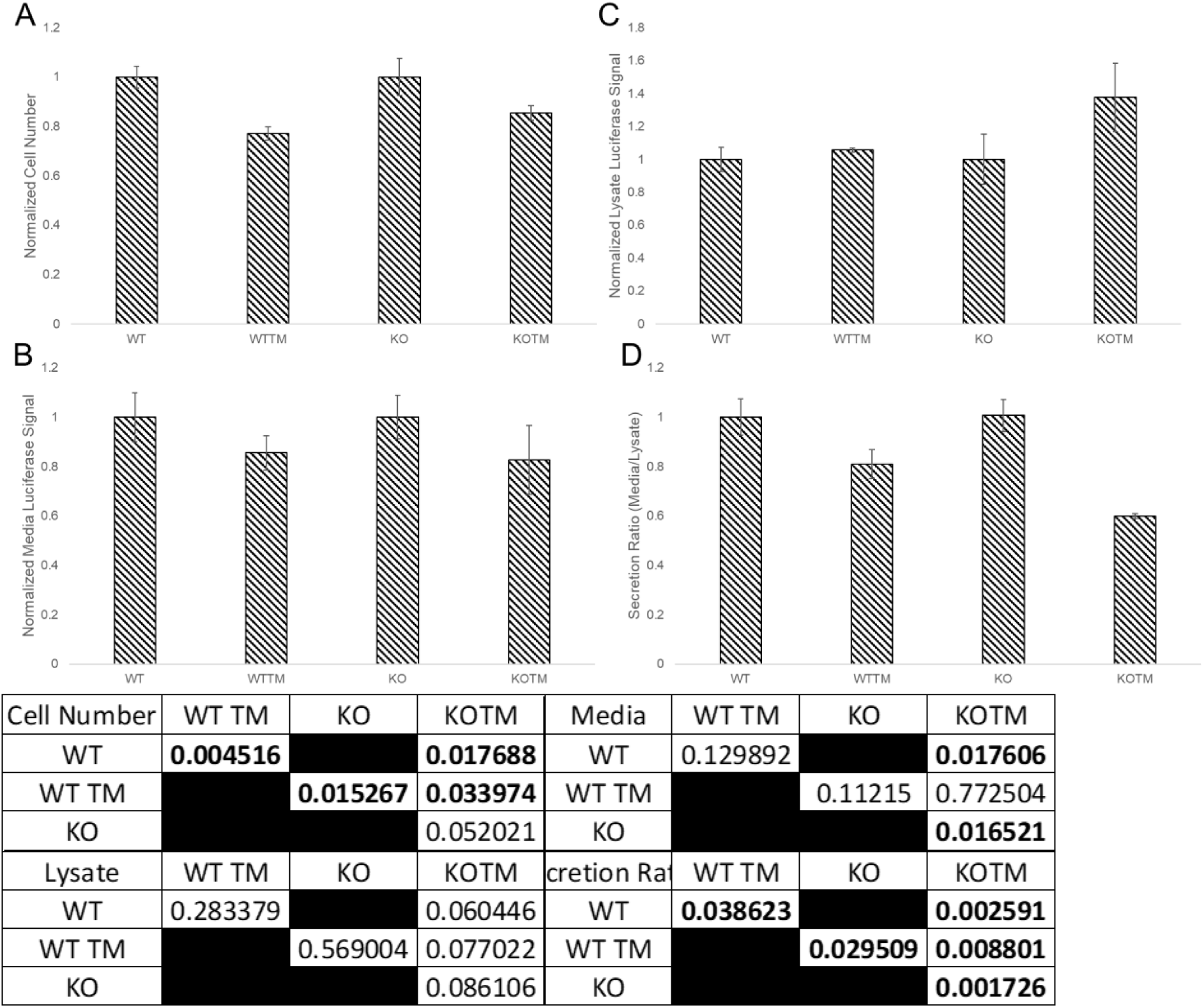
IRE1α knockout reduces secretion in response to TM. A) Normalized cell numbers for MDA-MB-231 cells that are wild type (WT), wild type with 5 µg/mL TM (WTTM), IRE1α^-/-^ (KO), and IRE1α^-/-^ with 5 µg/mL TM (KOTM) 24 hours after transfection of Gluc mRNA. B) Normalized 24-hour media Gluc accumulation for WT, WTTM, KO, and KOTM. C) Normalized Gluc in cell lysate for WT, WTTM, KO, and KOTM. D) Secretion ratio for WT, WTTM, KO, and KOTM. *Error bars represent standard deviations. Table represents Student’s t-test pair-wise p-values, N = 3. Bold indicates p < 0.05*.

### Autophagy inhibition improves secretion ratio

Under ER stress, in addition to IRE1α activation of XBP1-s transcription, in some cases autophagy is activated as a pro-survival mechanism(Das et al. 2020). Having determined that was the case here (Supplemental Figures 2A-L) we investigated how autophagy activation affected Gluc secretion. We used the class III phosphoinositide 3-kinase (PI3K) inhibitor, 3-methyladenine (3MA), to prevent the formation of phagophores and inhibit autophagy. The treatments of TM, 3MA, and both (3MATM) did not significantly alter the number of HeLa cells. However, the amount of Gluc secreted into the media was strongly correlated with cell number (Figure 3A), with 3MA increasing cell number and protein secreted both in the absence and presence of TM. The amount of Gluc in the cell lysate was not affected by exposure to TM or cotreatment of 3MATM. A plot of cell number vs lysate showed weak correlation, with the slope not statistically different from zero (Figure 3B). 3MA treatment alone caused no change in the secretion ratio of Gluc. In contrast, autophagy inhibition in combination with TM exposure increased secretion over TM alone, albeit lower than control (Figure 3C). Therefore, autophagy inhibition increased the likelihood of a synthesized Gluc protein being secreted rather than retained in the cell during ER stress. We confirmed the inhibition of autophagy and the corresponding changes in secretion using confocal microscopy (Supplemental Figures 1A-L).

**Figure 3.**
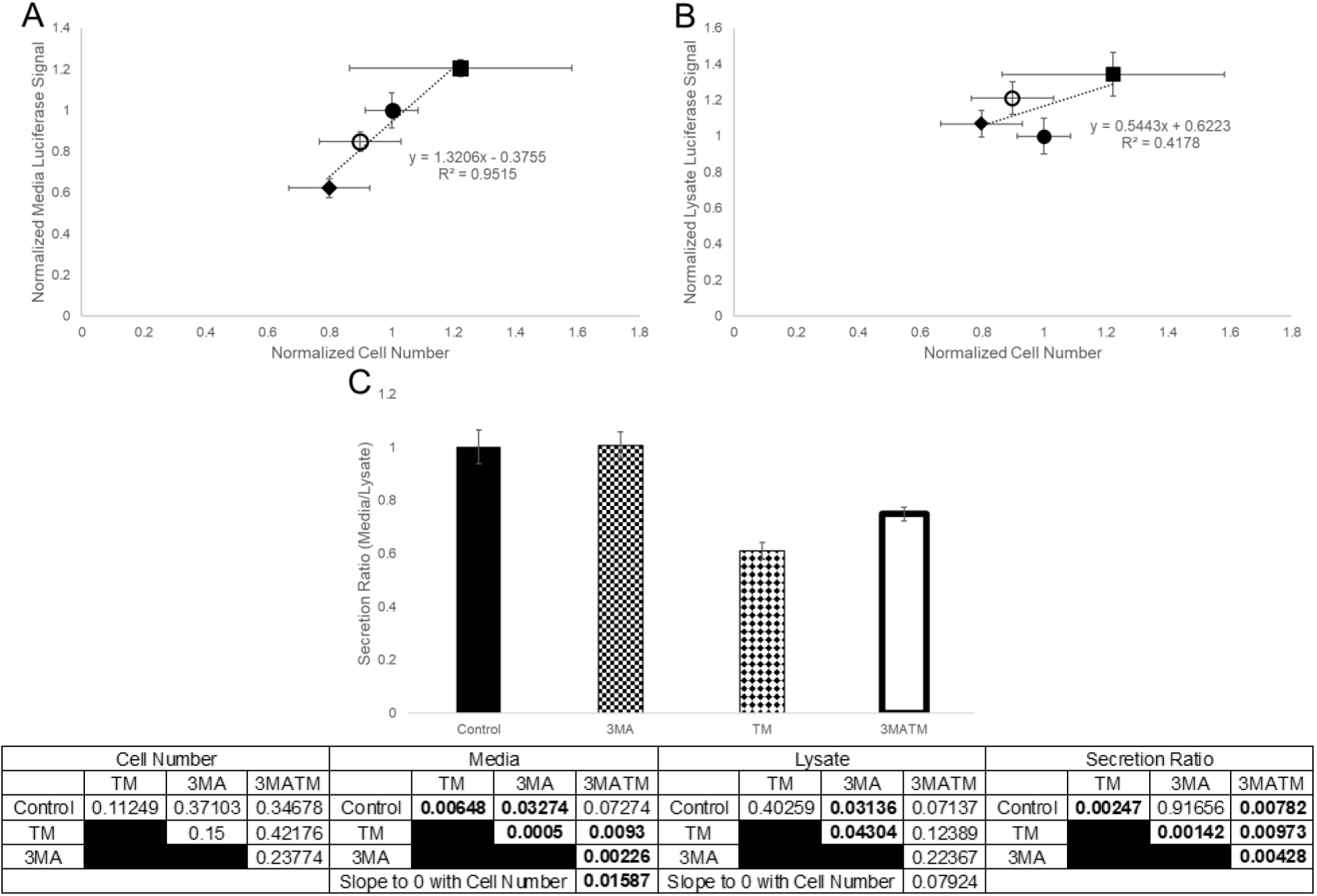
Effects of autophagy inhibition and TM-induced ER stress on secretion ratio. HeLa cells transfected with mRNA coding for Gluc and allowed to recover in control-(filled), 3MA-(squares), TM-(diamonds), or 3MA with TM-media (unfilled). A) Normalized 24-hour media accumulation of Gluc vs. normalized cell number. B) Normalized Gluc in cell lysate vs. normalized cell number. C) Secretion ratio of control, 3MA-treated, TM-treated, and 3MATM-treated cells. *The table is pairwise Student’s t-tests. Slope p-values calculated by linear regression and compared to a zero slope. Error bars represent standard deviation, N = 3. Bold indicates p < 0.05*.

### Proteasome activation restores secretion ratio

Autophagy and proteasomal degradation cooperate to address dysregulated proteostasis. To investigate how proteasomal degradation affected secretion of Gluc in our system, we used the proteasome inhibitor VR23 (VR) and proteasome activator (through PDE4 inhibition) Rolipram (RP). VR reduced the cell number and total Gluc secreted into the media for both TM-stressed and unstressed cells; cell number and Gluc secreted were correlated (Figure 4A). The amount of Gluc in the lysate increased with VR treatment for both the TM-stressed and unstressed cells (Figure 4B). Thus, VR decreased the secretion ratio, and co-treatment with TM compounded this effect (Figure 4C). Counterintuitively, proteasome inhibition, which should increase the amount of protein available for secretion, increased intracellular accumulation of Gluc and reduced secretion.

**Figure 4.**
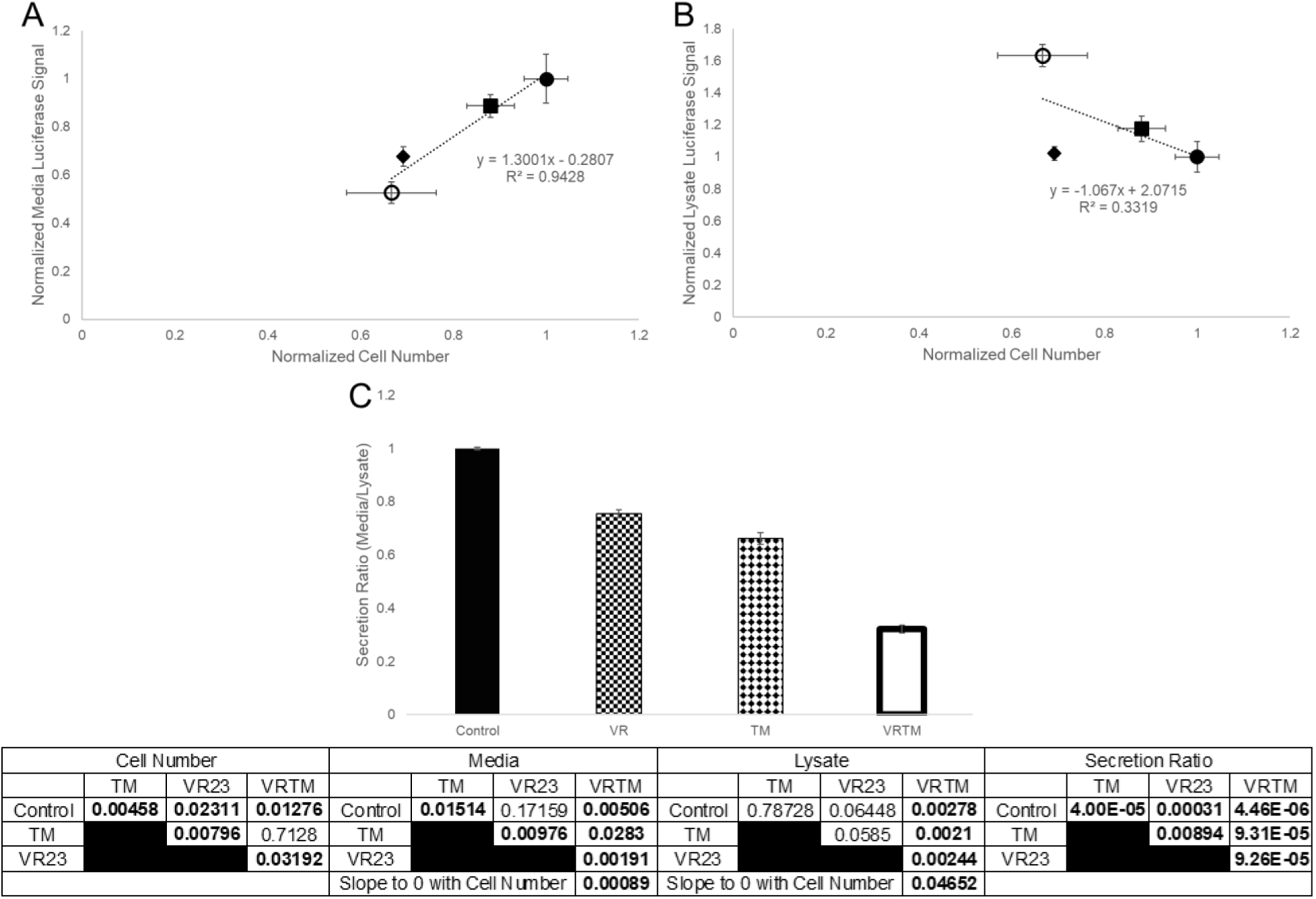
Proteasome inhibition reduced secretion. HeLa cells transfected with mRNA coding for Gluc and allowed to recover for 24 hours in control-(filled), VR23-(squares), TM-(diamonds), or VR23 with TM-media (unfilled). A) Normalized media accumulation of Gluc vs. normalized cell number. B) Normalized Gluc in cell lysate vs. normalized cell number. C) Secretion ratio of control cells and cells treated with VR23 alone, TM alone, and VR23 with TM. Slope p-values calculated by linear regression and compared to a zero slope. *The table is pairwise Student’s t-tests. Error bars represent standard deviation, N = 3. Bold indicates p < 0.05*.

To support this result, we examined whether proteasome activation would concomitantly increase secretion. Indeed, RP activation of proteasomes increased secretion in both the stressed and unstressed cells (Figure 5A), without altering the amount of Gluc in the lysates, regardless of TM treatment (Figure 5B). Thus, RP treatment increased the secretion ratio in both the TM-treated and unstressed cells (Figure 5C). Proteasomal activation was accompanied by a downregulation in autophagy (Supplemental Figures 1A-F and 1M-X), suggesting that sequestration of secreted proteins in cells occurs through autophagy.

**Figure 5.**
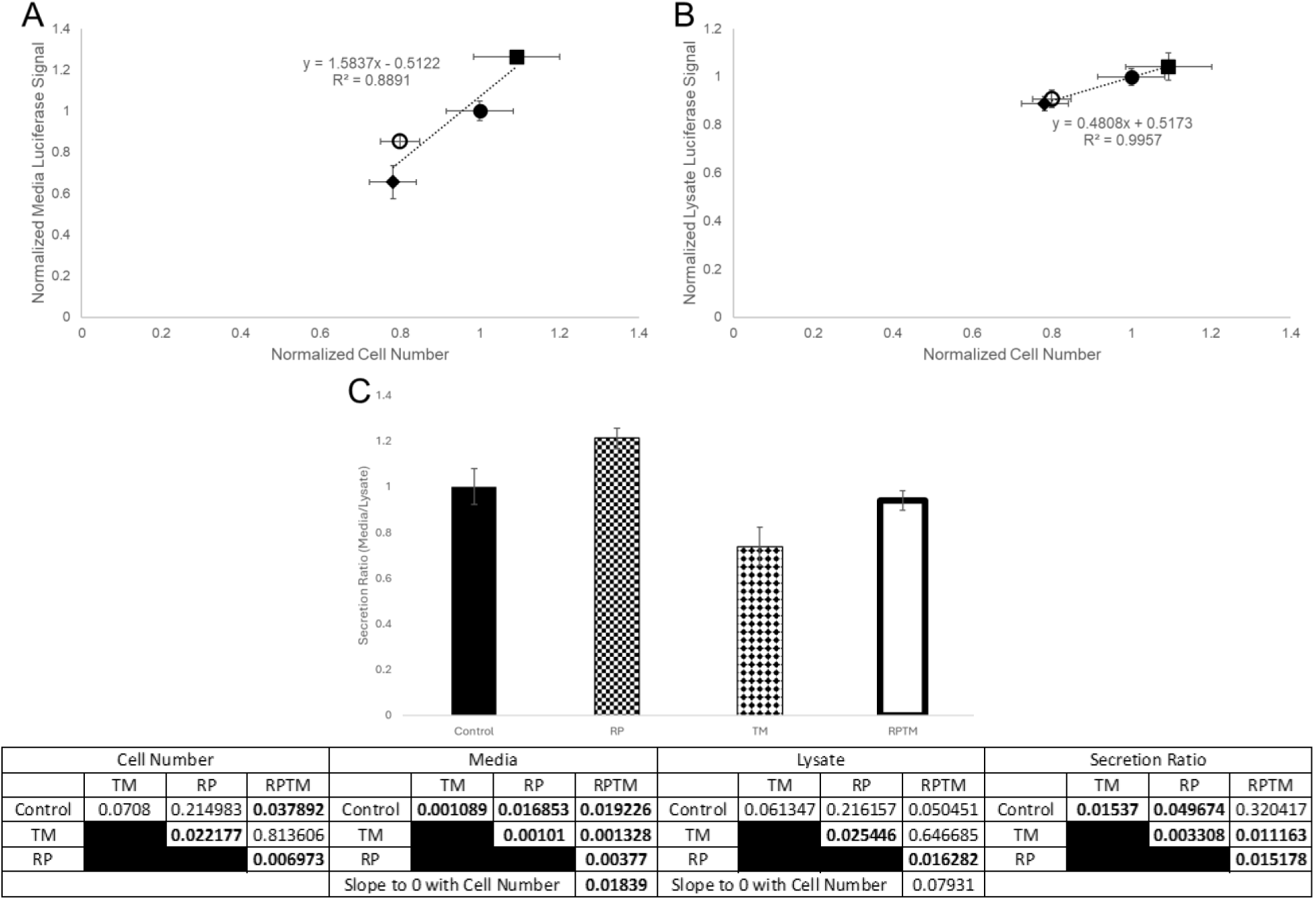
Proteasome activation improved secretion. HeLa cells transfected with mRNA coding for Gluc and allowed to recover for 24 hours in control-(filled), RP-(squares), TM-(diamonds), or RP with TM-media (unfilled). A) Normalized media accumulation of Gluc vs. normalized cell number. B) Normalized Gluc in cell lysate vs. normalized cell number. C) Secretion ratio of control cells and cells treated with RP alone, TM alone, and RP with TM. Slope p-values calculated by linear regression and compared to a zero slope. *The table is pairwise Student’s t-tests. Error bars represent standard deviation, N = 3. Bold indicates p < 0.05*.

### Secretion ratio changes with RP treatment are independent of transfected nucleic acid

ER stress alters several translation pathways that could alter the expression of secreted proteins(Yoshida 2007). To determine if the changes in secretion ratio were due to transcriptional regulation, we transfected cells with mRNA and pDNA complexed with Lipofectamine 2000 (LF2K) (Figure 6A). Post-transfection treatment with TM, RP, RPTM, VR, and VRTM all showed the same trend on secretion ratio comparing mRNA- and pDNA-based protein secretion. 3MA and 3MATM treatment reduced pDNA-based protein secretion as compared to control and TM treatment respectively, whereas 3MATM improved mRNA-based protein secretion as compared to TM treatment (Figure 3). Overall, the transfections of mRNA hindered cell growth less than pDNA transfection (Figure 6B) with 3MA causing similar toxicity as TM in pDNA transfected cells, which could suggest pDNA transfection caused additional stress/cytotoxicity through increased LF2K concentration that autophagy helps to alleviate.

**Figure 6.**
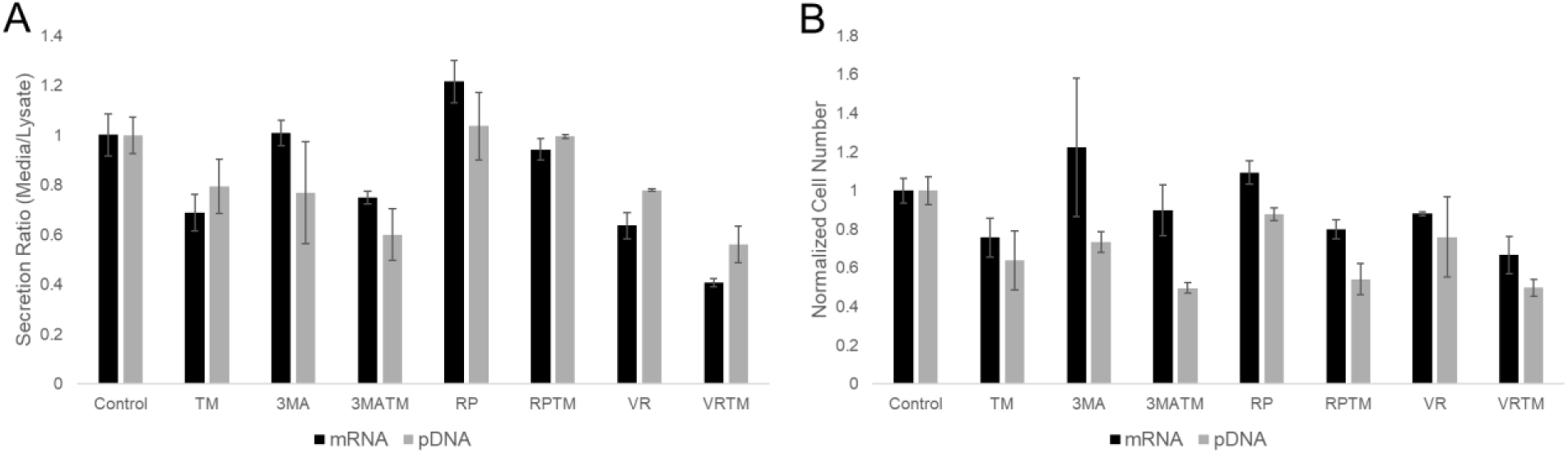
Changes to proteasomal degradation were independent of transfected nucleic acid type. A) Secretion ratio responses to small molecule alteration of degradation pathways with either mRNA (black) based protein expression or pDNA (gray) based protein expression. B) Relative cell number for each small molecule alteration of degradation and mRNA (black) or pDNA (gray) transfection. *Error bars represent standard deviations, N=3*.

These results confirmed that the effects of proteasomal activation/inhibition on secretion are due to alterations in translation and protein processing, not transcription.

### ER stress dependence on autophagy and proteasomal activity is cell type dependent

We used MDA-MB-231 and HeLa cells to determine if the influence of 3MA and RP on the secretion patterns under ER stress is cell type dependent. For MDA-MB-231 breast cancer cells, TM reduced secretion ratio (Figures 7A-C). Reducing degradation via 3MA and VR23 also reduced secretion ratio (Figures 7A and B), while increasing proteasomal degradation with RP did not statistically impact secretion (Figure 7C). The combination of TM and degradation inhibition further reduced secretion, relative to TM alone (Figures 7A and B). TM and RP treatment in MDA-MB-231 was not statistically different from TM alone (Figure 7C). The normalized number of cells remaining after 24 hours of protein production was lower for the MDA-MB-231 cells than HeLa cells, suggesting that apoptosis was more likely to be induced in MDA-MB-231 cells vs. HeLa cells under these conditions (Figure 7D). These results indicate that altering degradation processes during ER stress to increase protein production will be cell line dependent. Nonetheless, increasing proteasomal degradation either improves secretion (HeLa) or has no negative impact on secretion (MDA-MB-231), making it a consideration for protein production processes.

**Figure 7.**
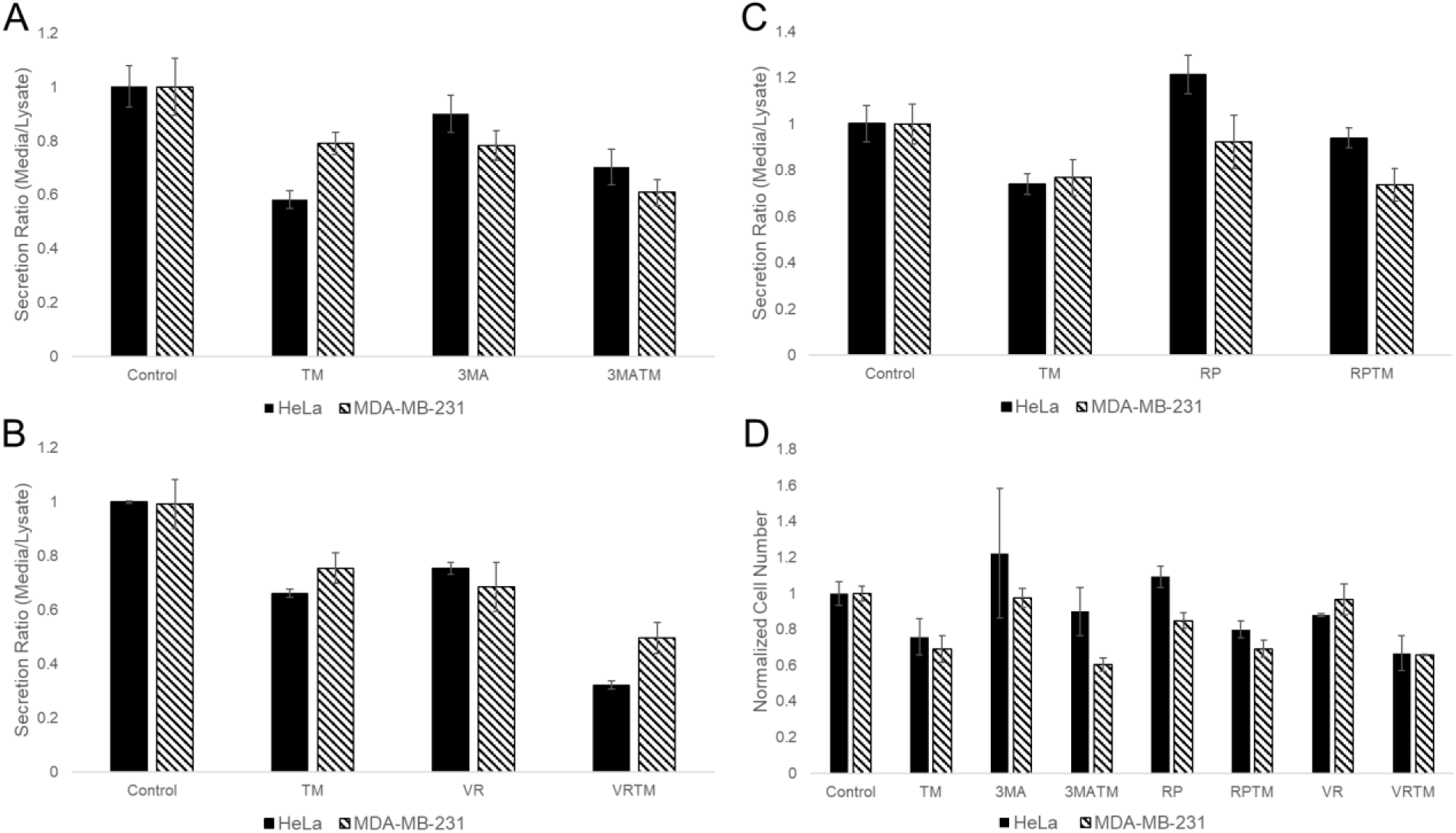
Restoration of secretion patterns altered by ER stress is cell line dependent. HeLa (solid) and MDA-MB-231 (striped) cells transfected with mRNA coding for Gluc and allowed to recover for 24 hours in media containing A) no small molecules, TM, 3MA, or both; B) no small molecules, TM, VR23, or both; and C) no small molecules, TM, RP, or both. D) Normalized cell number for each treatment (untreated, 3MA, RP, and VR23) and the cotreatments with TM.

## Discussion

In this work, we demonstrated that ER stress reduced the amount of Gluc secreted into the media not only by reducing cell number but also by reducing the fraction of synthesized proteins that are secreted rather than retained in the cells. Autophagy inhibition improved secretion, while, counterintuitively, increasing proteasomal degradation increased the likelihood of secretion. Thus, the capacity of cells to secrete synthesized proteins is related to the activity of its protein degradation mechanisms.

We focused this work on understanding how ER stress and the UPR influence protein secretion because, as stated above, cells used in industrial bioprocesses continually experience ER stress due to a variety of causes, which, in turn, reduces yield of expressed proteins. Our model for ER stress induction, administration of TM, is well-established for studying ER stress and the UPR and causes similar changes to those experienced by cells in bioprocesses, despite the differences in the laboratory and industrial contexts (e.g., chemical vs. physical stress)(Gerlach et al. 2012). For instance, nutrient and oxygen deprivation, heat stress, and shear stress disrupt N-glycosylation similar to TM(Gerlach et al. 2012).

Reduced protein yields can result from many bottlenecks. It is well-established that ER stress reduces cell growth and protein production(Savage and Baur 1983), consistent with our results here. But, the effects of ER stress on intracellular protein processing have not previously been explored(Ha et al. 2019). Disruptions in glycosylation impair the function of many transmembrane glycoproteins that control vesicle trafficking(Olden et al. 1979), making it reasonable to presume that ER stress would alter protein secretion. Our results here provide clear evidence that controlling intracellular processing of expressed proteins can directly improve the secretion of active protein.

ER stress upregulates the autophagic degradation pathways and ERAD as a pro-survival mechanism(Meusser et al. 2005; Ichimiya et al. 2020). These two forms of degradation play a cooperative role in decreasing the misfolded protein load that accumulates and leads to ER stress. Previous work indicated that induction of autophagy caused an increase in protein secreted into the media(Cavalli and Cenci 2020). However, they did not determine the per cell production or secretion ratio, only that the induction of autophagy is pro-survival and increases the number of cells producing protein. Our results showed that TM-induced autophagy hindered protein production, when normalized for cell number, and that inhibiting TM-induced autophagy improved protein secretion per cell. This suggests that there is an optimal level of autophagy that results in the highest protein production per cell with the optimum survivability of cells under ER stress.

Since autophagy inhibition improves secretion and proteasomal degradation could decrease autophagy, we investigated RP to increase proteasomal degradation and ERAD. RP is a cAMP-specific phosphodiesterase 4 (PDE4) inhibitor that is used clinically to treat neurological disorders(Lokireddy et al. 2015). Increased cAMP levels increase proteasome phosphorylation and activity. RP treatment increases cell number and protein secretion without increasing protein accumulation in cells. The increase in protein secretion of cells treated with RP could be due to improved protein trafficking, since reduced ER stress would result from increased degradation of unfolded proteins. Additionally, translation could be generally accelerated due to the resulting increased concentration of free amino acids in the cytosol that result from degraded proteins.

Elevated cAMP levels could also activate other pathways, including the AMP-activated Protein Kinase pathway (Aslam and Ladilov 2022), that promote protein production and cell viability. We demonstrated that inhibition of proteasome activity reduced Gluc secretion per cell and increased intracellular Gluc accumulation and autophagy, thereby highlighting the importance of both autophagy and proteasomal degradation in maintaining cellular protein secretion capacity. We expect that our results will inform the future development of therapeutic protein bioprocesses, by encouraging investigation of methods to increase misfolded protein degradation without increasing autophagy.

## Author Contributions

RCS - methodology, investigation, data acquisition, conceptualization, writing – original draft, reviewing and editing, visualization. CC - methodology, conceptualization, writing - reviewing and editing. SPW - methodology, conceptualization, writing - reviewing and editing.

## Acknowledgements

We would like to thank the members of the Cellular and Biomolecular Laboratory for their input and advice on this project, especially Kevin Chen for providing the MDA-MB-231 wild type and IRE1α-/- cells and Courtney Bair for assisting with cell culture. We would like to thank Dr. Daniel Vocelle for assistance with the Attune Cytpix Flow Cytometer, located in the MSU Flow Cytometry Core Facility, supported by the Equipment Grants Program, award #2022-70410-38419, from the U.S. Department of Agriculture (USDA), National Institute of Food and Agriculture (NIFA). Finally, we would like to thank Dr. Melinda Frame at the Center for Advanced Microscopy at MSU for assisting with Confocal Laser Scanning Microscopy.

## Abbreviations

3-methyladenine - 3MA

3-methyladenine and tunicamycin - 3MATM

Activating Transcription Factor 4 - ATF4

Activating Transcription Factor 6 - ATF6

C/EBP homologous protein - CHOP

Chaperone Mediated Autophagy - CMA

Endoplasmic Reticulum - ER

ER DnaJ 3 - Erdj3

ER DnaJ 4 - Erdj4

ER DnaJ 5 - Erdj5

ER-associated degradation - ERAD

eukaryotic translation initiating factor 2α - eIF2α

*Gaussia princeps* Luciferase - Gluc

Heat Shock Protein A8 - HSPA8

Inositol-Requiring Enzyme 1α - IRE1α

IRE1α^-/-^ Knockout - KO

IRE1α^-/-^ Knockout with tunicamycin - KOTM

Lipofectamine 2000 - LF2K

Microtubule-associated proteins 1A/1B light chain 3B - LC3B

Protein Kinase R-like Endoplasmic Reticulum Kinase - PERK

Rolipram - RP

Rolipram with tunicamycin - RPTM

Site-1 protease - S1P

Site-2 protease - S2P

Suppressor Enhancer Lin-12-Like - SEL1L

Tunicamycin - TM

Ubiquitin Protein Ligase E3 Component N-Recognin 4 - UBR4

Ubiquitin Protein Ligase E3 Component N-Recognin 5 - UBR5

Unc-51-like Kinase 1 - ULK1

Unfolded Protein Response - UPR

VR23 - VR

VR23 and tunicamycin - VRTM

Wild Type - WT

Wild Type with tunicamycin - WTTM

X-box Binding Protein-1 - XBP1

XPB1-spliced - XBP1-s

