## Supplemental Figures for "Secreted Protein Production is Improved by Controlling Endoplasmic Reticulum Stress Associated Protein Degradation"

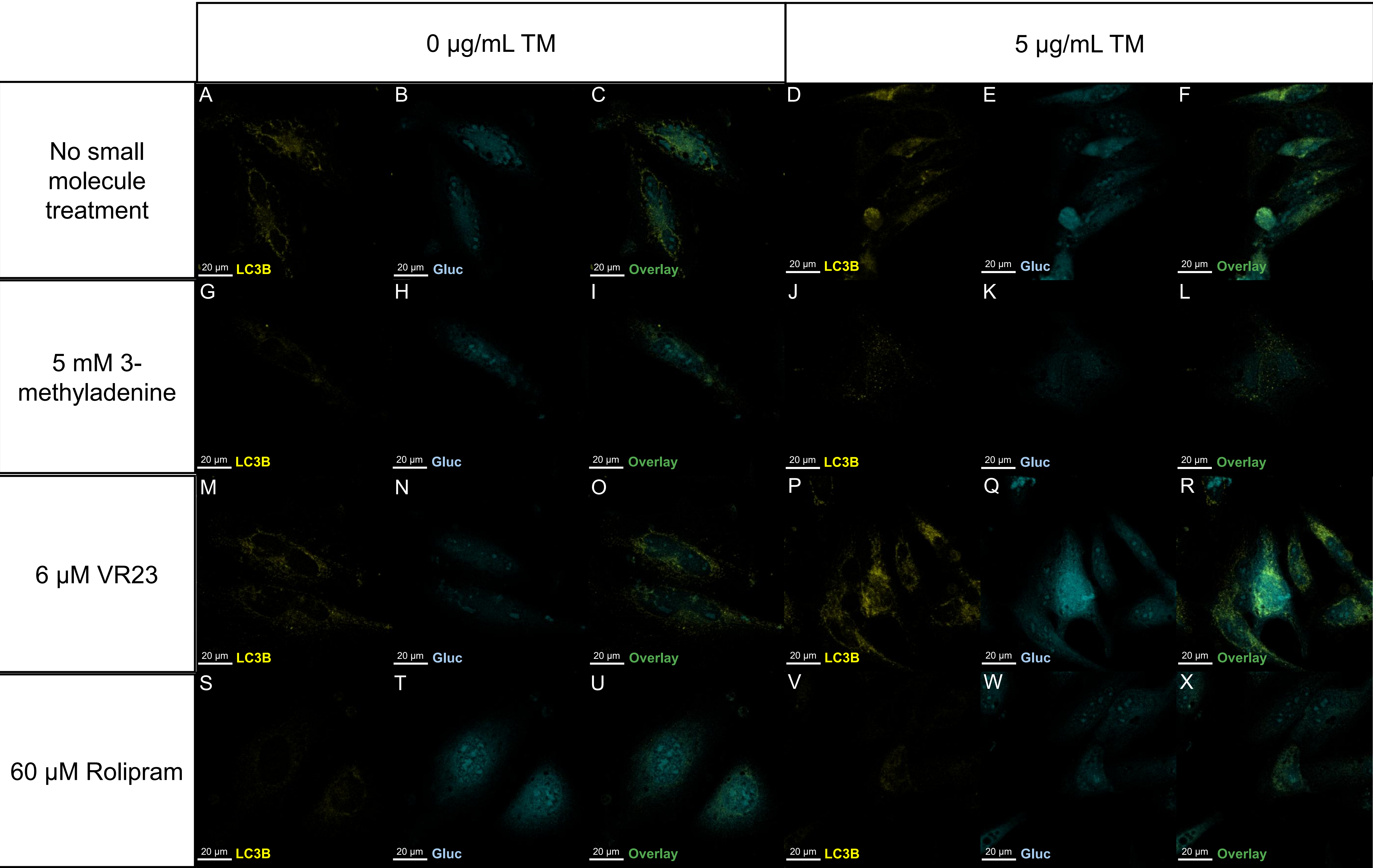
0 µg/mL TM
5 µg/mL TM
No small molecule treatment
5 mM 3-methyladenine
6 µM VR23
60 µM Rolipram
A
B
C
D
E
F
G
H
I
J
K
L
M
N
O
P
Q
R
S
T
U
V
W
X
20 µm
LC3B
Gluc
Overlay

**Supplemental Figure 1** Micrographs of HeLa cells taken 48 hours post transfection with pDNA encoding CFP-Gluc and allowed to recover in the presence or absence of TM and small molecules affecting degradation. (A,D,G,J,M,P,S,V) LC3B-yellow; (B,E,H,K,N,Q,T,W) Gluc-blue; and (C,F,I,L,O,R,U,X) the overlay of LC3B and Gluc. A-C) no small molecule drugs, D-F) 5 µg/mL TM, G-I) 5 mM 3MA, J-L) 5 µg/mL TM and 5 mM 3MA, M-O) 6 µM VR23, P-R) 6 µM VR23 and 5 µg/mL TM, S-U) 60 µM RP, and V-X) 60 µM RP and 5 µg/mL TM.

Comparing micrographs of control cells (SF1A-C) with TM treated cells (SF1D-F) showed that in each cell more LC3B autophagy marker and Gluc are retained. 3MA treatment inhibited autophagy as shown by the lack of LC3B in both TM-stressed and unstressed cells (SF1G and J) with a reduction in intracellular Gluc signal (SF1H and K). VR23 treatment caused an increase in autophagy (SF1M). VR23 and TM cotreatment caused the most significant autophagy increase and Gluc accumulation (SF1P-R). RP treatment reduced the LC3B signal (SF1S and V) without impacting Gluc signal (SF1T and W). These findings are consistent with data generated by cell lysate luciferase activity assays.

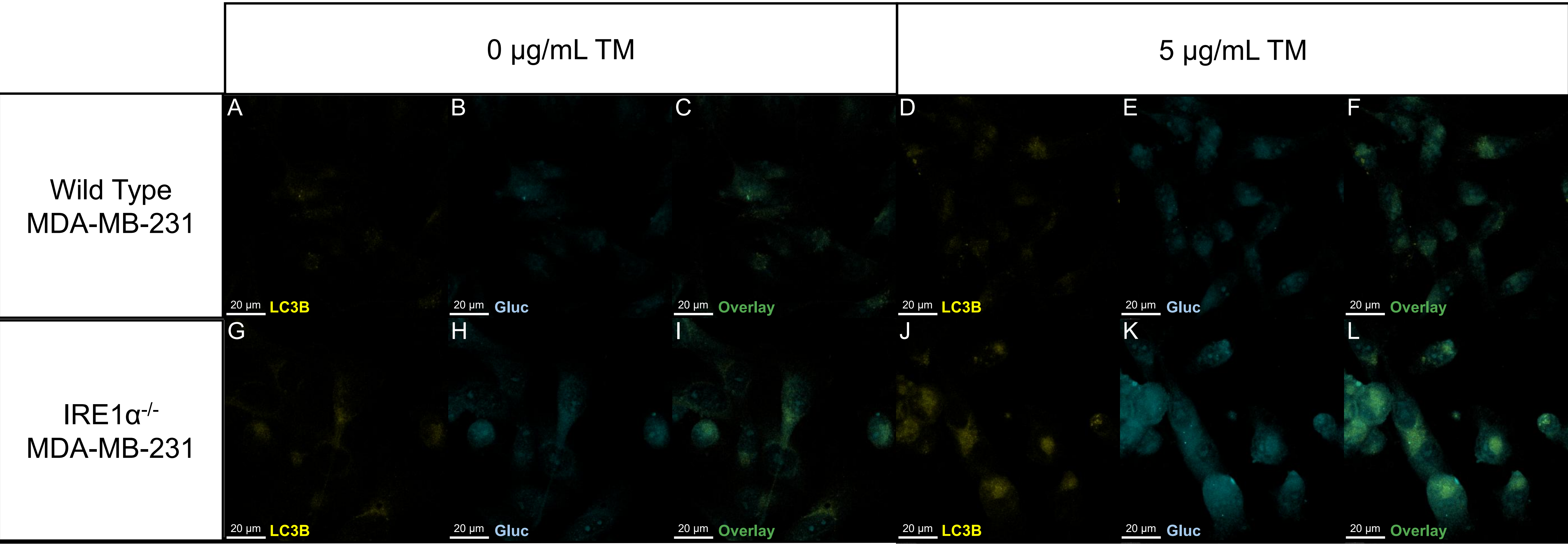
0 μg/mL TM
5 μg/mL TM
Wild Type MDA-MB-231
IRE1α-/- MDA-MB-231
A
B
C
D
E
F
G
H
I
J
K
L
20 μm
LC3B
Gluc
Overlay

**Supplemental Figure 2** Micrographs of MDA-MB-231 WT (A-F) and KO (G-L) cells taken 48 hours post transfection with pDNA encoding CFP-Gluc and allowed to recover in the absence (A-C and G-I) or presence (D-F and J-L) of TM.

TM treatment caused an increase in the LC3B signal and Gluc signal in both the WT and IRE1α-/- cells (compare SF2A/B with D/E and G/H with J/I). TM increased retention of both LC3B and Gluc in the IRE1α-/- cells as compared with control (compare SFD/E with J/I). These findings are consistent with cell lysate luciferase activity measurements.
